# Dynamics of Excitability in Axonal Trees

**DOI:** 10.1101/2025.07.24.666461

**Authors:** Laurie D. Cohen, Tamar Galateanu, Shimon Marom

## Abstract

We report that axons of cortical neurons, structurally intricate excitable media, maintain remarkably high fidelity in transmitting somatic spike timing, even during complex spontaneous network activity that includes extremely short (2–3 msec) inter-spike intervals. This robustness underscores their function as reliable conducting devices under physiological conditions. It is nevertheless well established that under artificially imposed, high-rate pulsing stimuli, axonal conduction can fail, with vulnerability depending on distance and branching. In line with this, we demonstrate that conduction failures can also occur at frequencies as low as 10 Hz, provided that stimulation is sustained for several seconds. Under these conditions, propagation delays increase and failures accumulate, particularly in distal branches, whereas effects are negligible at 1–4 Hz. Simulations incorporating cumulative sodium channel inactivation at vulnerable sites reproduce these dynamics. Our findings refine the view of axons as active, heterogeneous structures: they are exceptionally reliable across most physiological regimes, yet exhibit limits under prolonged or extreme stimulation, a regime rarely encountered *in vivo* but critical for understanding axonal excitability.

**Significance Statement:** Axons are widely regarded as reliable conduits of action potentials from the neuron cell body to downstream targets. We confirm this reliability during spontaneous network activity, even when spikes occur in complex patterns with very short interspike intervals. Prior studies have shown that conduction can fail under rapid, high-frequency trains of artificial stimuli. Here we extend this principle by showing that sustained stimulation at 10 Hz—a frequency within physiological ranges—can also induce propagation delays and failures, particularly in distal and higher-order axonal branches. Simulations incorporating cumulative sodium channel inactivation reproduce these effects. Thus, axons should be understood as highly reliable under natural conditions, yet context-dependent, with excitability shaped by the history of sustained activity.

## Introduction

In 1965, George H. Bishop published a beautiful account on his scientific “life among the axons,” where he wrote “By tacit general consent so far, the axon does not think. It only ax.”^1^ Over the past sixty years, a rich body of knowledge has accumulated, demonstrating a range of parameters that influence spike propagation through axons. The picture that emerges (e.g.^2–14^) is that action potentials are generated in the axon initial segment, a complex anatomical site with specialized, adaptive protein machinery. Once generated, the action potential travels through a kinetically adaptive and structurally heterogeneous tree. The short- and long-term plasticity of excitability matters; the topology of the axon as well as its local dimensions and microstructure matter; differential distribution of ionic channels along the axon is critical, as is the rich repertoire of gating processes of ionic channels; analog variations in sub-threshold membrane potential are important sources of adaptive dynamics of excitability; mechanical and thermal changes matter; and more.

Progress in understanding the functional consequences of axonal inhomogeneities on excitability depends on two sources of data: The first comes “from below”, that is, exposing unique microscopic axonal elements and their dynamics. For instance, revealing the distributions and densities of different ionic channels along axons,^4,15–17^ or the protein content of varicosities,^4,18,19^ or the ratios between diameters of axons at branching points,^4,12,20^ etc. These data are invaluable, as they provide the necessary pieces to construct understanding. The second comes “from above”, the sum impact of the microscopic infrastructure on key functional axonal excitability measures, as spikes invade deeper into the axonal tree: axonal refractoriness, propagation velocity, variations in time of arrival to points along the axon, and probability of action potential propagation failure.

The present report describes an experimental study that falls under the latter category, offering a macroscopic view of axonal excitability. The study specifically focuses on the dynamic changes in excitability within deep axonal trees. The key factors considered are the physical distance from the spike source to a specific point in the axon, and the tree depth, which is the number of branching points counted from the source to that point. To achieve this, we employed direct electrical recording of individual spikes as they propagate down the axonal tree of a cortical neuron. The *in-vitro* approach was chosen due to the challenges associated with conducting high spatial resolution direct electrical recording. We presume that the biophysical nature of the questions discussed here is sufficiently broad to transcend the distinctions between *in-vivo* and *in-vitro* conditions.

Two main results are described: (1) When spikes are naturally ignited by network activity, dynamics of excitability deep in the neuronal axonal tree do not constrain reliable spike propagation. The time delay between any pair of spikes detected in the cell body is reliably maintained while the pair of spikes is conducted down the axonal tree. (2) In contrast, direct, noninvasive stimulation of a neuron within a physiologically relevant frequency – as low as 10 Hz if sustained for several seconds or longer – gives rise to time-dependent and frequency-dependent changes in spike propagation delays and spike propagation failures. These effects monotonically increase as a function of distance and branching points along the way, whereas they are negligible at 1-4 Hz. The observation is interpreted as reflecting increased sensitivity to propagation failure at branching points due to slow, cumulative inactivation of axonal sodium channels.^21–25^

## Materials and Methods

### Animal welfare

Experiments were performed in primary cultures of newborn rat neurons prepared according to a protocol approved by the Technion, Israel Institute of Technology Committee for the Supervision of Animal Experiments (Approval IL-159-12-24).

### Primary cultures

of rat cortical neurons were prepared as described previously^26^. Briefly, cortices of 0-to 1-day-old rats (Wistar, either sex; Charles River Laboratories) were dissected, dissociated by trypsin treatment (T1426; Merck-Sigma Aldrich) and DNase (10104159001; Merck-Sigma Aldrich), followed by trituration using a siliconized Pasteur pipette. The cells (150 × 10^3^) were plated on HD-MEA arrays (MaxOne Single-Well Planar Microelectrode Arrays; MaxWell Biosystems). Before cell plating, the arrays were sterilized for 30 minutes in 70% ethanol, rinsed with tissue-culture grade water, and coated with polyethylenimine (P3143; Merck-Sigma Aldrich). Cells were initially grown in medium containing Minimum Essential Medium (M2279; Merck-Sigma Aldrich), 25 mg/l insulin (E-5500; Merck-Sigma Aldrich), 20 mM glucose (Merck-Sigma Aldrich), 2 mM L-glutamine (Merck-Sigma Aldrich), and 10% Nu-Serum (FAL355100, Lapidot Pharma or Becton Dickinson Labware). The preparation was then transferred to a humidified tissue culture incubator and maintained at 37°C in a 95% air and 5% CO_2_ mixture. Starting seven days after plating, half the volume of the culture medium was replaced three times a week with cell culture medium like the initial medium but devoid of Nu-Serum, with a lower concentration of L-glutamine (0.5 mM), and supplemented with NS21 (50X, P07-20021; Pan Biotech). Experiments were conducted 3 to 4 weeks after plating. For experiments in isolated cortical neurons, network reverberations were minimized by bath application of synaptic blockers (D,L-2-amino-5-phosphonopentanoic acid (APV, 100 µM; Merck-Sigma Aldrich), 6-cyano-7-nitroquinoxaline-2,3-dione (CNQX, 10 µM; Merck-Sigma Aldrich), and 50 µM bicuculline methiodide (BIC; Merck-Sigma Aldrich). In networks where the synaptic blockers visibly did not exert the full effect, APV and BIC were applied at 5X of the nominal concentrations.

### Electrophysiology

Recordings were performed using the complementary-metal-oxide-semiconductor (CMOS)-based high-density microelectrode array (HD-MEA) technique (MaxWell Biosystems AG). The technique, developed over the past decade^27–30^, enables non-invasive measurement of electrical aspects related to action potential spreading. It practically imposes no constraints on the duration of recording sessions and allows access at good enough spatiotemporal resolution to arbors of cortical axons, including tiny axon terminal branches. The technique was validated by its developers in a series of experiments, showing that the footprint of elaborate axonal trees can be reconstructed from their electrical activities, nicely matching histological morphology^27,29,31^. Critical for the subject of the present work, the technique enables estimation of changes in spike times of arrival across axonal arbors. The HD-MEA device was installed in a 37°C CO_2_ incubator and (in our case) retrofitted with a customized Peltier cooling system that further protects against temperature variations due to heat produced when the chip is working. Signals were amplified, high pass filtered at 300 Hz, and digitized (10-bit, 20 kHz) on chip. In a typical experiment, we scanned neuronal units and their spontaneous activities and identified candidate axonal footprints (this was done with the MaxLab Live software environment, MaxWell Biosystems AG). Application Programming Interface (API) was used to select an electrode configuration encompassing a given neuron and to route selected electrodes for recording.

### Simulations

expanding a model of axonal propagation convolved with slow inactivation^32^ were run in NEURON^33^ version 8.2.6 called from Python 3.12. Here we constructed a long axon that branched into two daughter branches. A 10-Hz repeated activation at a 100-µm short section (“AIS”) that we positioned just before the axon provided the current for the evoked action potentials. Parameters were *C*m = 1 µF/cm^2^, *Ra* = 100 Ω·cm; gNa_max_ was varied between 65-120 mS/cm^2^, gK_max_ = 36 mS/cm^2^, gleak = 0.3 mS/cm^2^, E-leak = −54.3 mV, *T* = 20°C. Spatial discretization was based on d-lambda rule. The geometric ratio (GR) was calculated as in^12^.

## Results

We begin our description of the results by demonstrating that pairs of spontaneously evoked spikes, limited by somatic refractoriness and primarily resulting from ongoing network activity, persistently maintain the precise inter-spike interval throughout the axonal tree. This suggests that, at least in the context of spontaneous activity, axonal refractoriness does not impose limitations on the high temporal resolution dictated by somatic refractoriness.

Traditionally, single neuron refractory period is directly estimated using a double pulse protocol applied to the membrane of the neuron. This method allows for the identification of absolute, relative, and in some cases, supernormal refractory periods.^10^ The approach used here involves studying axonal refractory period by taking advantage of the rich temporal structure of spontaneous activity in cortical networks.^26,34,35^ When long enough stretches of spontaneous data are collected, one can identify occurrences of two somatic spikes appearing in close temporal proximity (<7 msec), enabling examination of the extent to which the cell-body inter-spike interval is maintained down the axonal tree.

Figure 1 presents an axonal tree object, constructed offline utilizing a minimum spanning algorithm (Wolfram *Mathematica* environment) that connects axonal electrodes associated with a specific neuron. The list of axonal electrodes was retrieved from the data generated by MaxLab Live software (MaxWell Biosystems). An electrode depicted as ‘source’ signifies a soma-related position, likely representing the axon initial segment. Axonal electrodes exhibiting an average signal-to-noise ratio exceeding 4.5 are colored based on their time of arrival (TOA): the time elapsed between the source’s spike appearance and its detection at each axonal electrode. Letter-designated arrows point to several axonal electrode positions; superposed traces recorded with these electrodes are shown on the right panel of Figure 1, positioned along a time axis (see bottom of figure) according to the time of arrival. Although the absolute amplitudes within each set of traces vary substantially due to low-frequency components (‘noise’), the axonal spikes are clearly detectable, and the TOA seems consistent. In our experimental setup, the estimated average axonal propagation velocity, derived from 9557 axonal segments (corresponding to 560 neurons), is 0.5 m/s (standard deviation 0.12), commensurate with previous findings.^27,29^.

**Figure 1:**
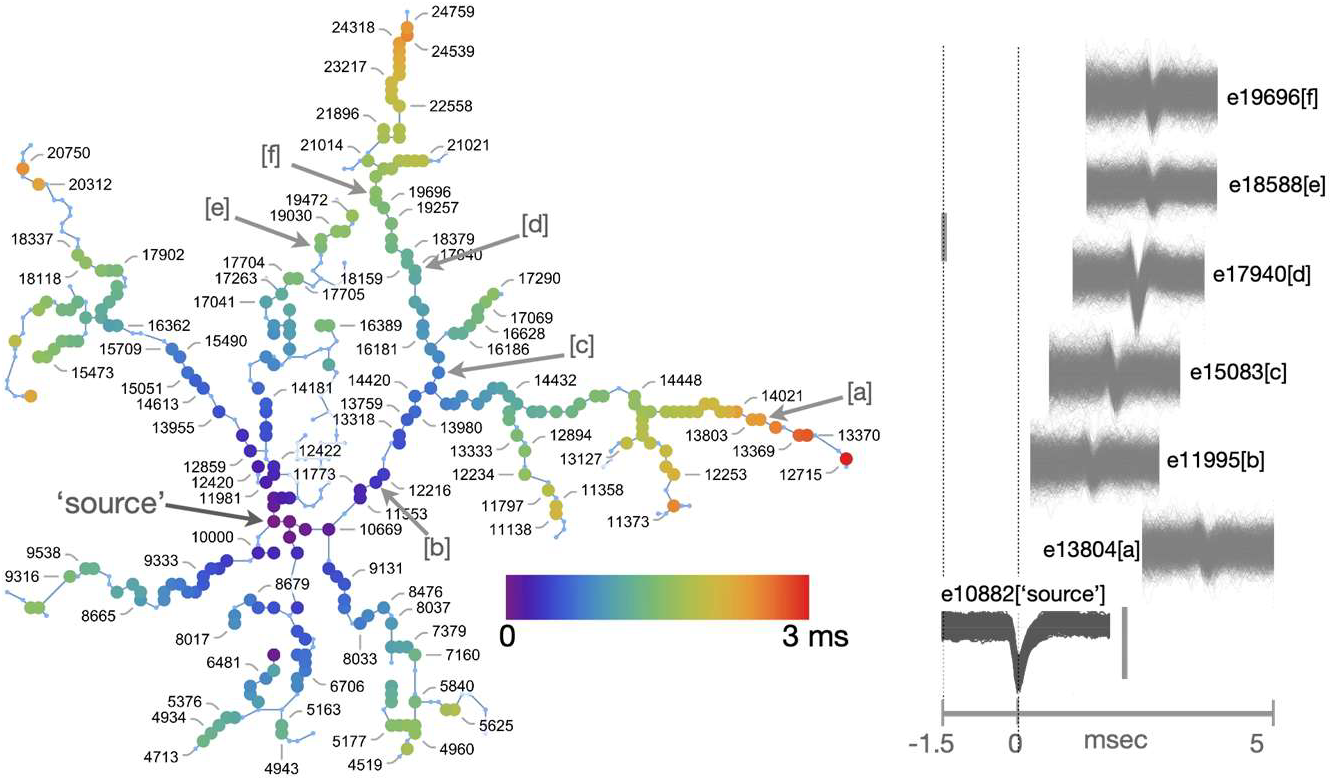
Electrophysiological recordings from axonal branches. (**Left**) An axonal tree object is constructed offline using a minimum spanning algorithm (Wolfram Mathematica environment) that connects axonal electrodes associated with a specific neuron. The HD-MEA electrodes underneath axon branches of a single neuron are retrieved from the output of MaxLab Live software (MaxWell Biosystems AG), an unsupervised object tracking algorithm that detects the propagating action potential signals and follows the path traveled from the initiation site to distal branches. Axonal electrodes exhibiting an average signal-to-noise ratio exceeding 4.5 are color-coded, indicating time of arrival (TOA, msec) relative to spike time recorded by an electrode at the ‘source’ (soma-related position, marked with a black arrow, likely representing the axon initial segment). Letter-designated arrows [a-f] indicate several axonal electrode positions. (**Right**) Hundreds of signals (superposed) recorded simultaneously at the arrow-depicted electrodes, temporally aligned along a time-axis according to the time of arrivals relative to the source spike events (bottom trace). Mean spike amplitudes: axonal traces: e19696: 6 µV, e18588: 4 µV, e17940: 12.5 µV, e15083: 5.3 µV, e11995: 3.55 µV, e13804: 3.3 µV. ‘Source’ traces: 122 µV. Scale bar: axonal traces: 20 µV; ‘source’ traces: 150 µV.

To quantify the reliability of maintaining spike intervals along axonal trees, we extracted traces recorded at the source, in which two spikes appear in close temporal proximity. Figure 2A (top) shows a group of 97 superposed source traces extracted from the data according to a strict criterion: they all must include two spikes with peaks that cross a -62.6 µV (10 LSB) threshold and a time difference smaller than 7 msec. The bottom group shows the corresponding traces recorded in one of the loci deep inside the axonal tree. The time delay between the first spikes of the source and the axon is 2 msec.

**Figure 2:**
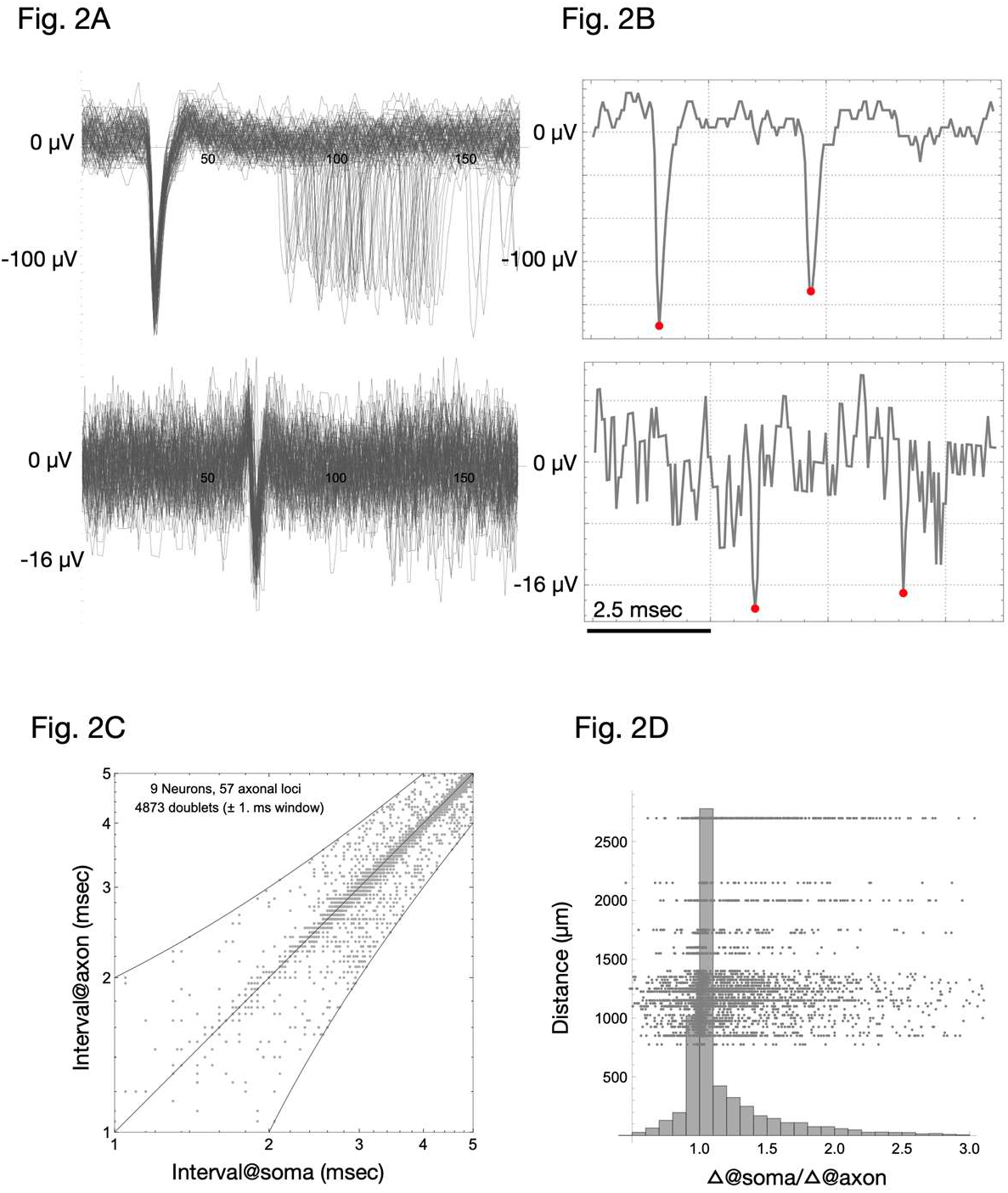
Reliability of maintaining spike intervals along axonal trees. (**A**) Spike traces were extracted from the recordings at the source according to a strict criterion: they must include 2 spikes with peaks that exceed a 10 µV threshold and a time difference smaller than 7 msec. Top panel: 97 superposed pairs of spikes accrued in the source electrode matching the described criteria. Bottom panel: the corresponding traces in the selected axonal electrode, at one of the loci deep inside the axonal tree (1350.3 µm calculated shortest path from the source). There is a 2 msec time delay between the first spike of the source and the first spike at the axon. (**B**) Example of one somatic trace and its corresponding axonal trace. A single trace (one of the 97 of panel A, same electrodes) from the ‘source’ electrode (top) and axonal electrode (bottom). The positions of the two spikes (estimated by an algorithm described in the main text) are marked in red. (**C**) Relations between the inter spike interval in the axonal electrode (ordinate) and soma electrode (abscissa); data from 9 neurons, 57 axonal loci, total of 4783 spike doublets (±1 msec window). Neurons are from four network preparations. Most of the points concentrate around the unity line. Data are shown on logarithmic scales. (**D**) Histogram of the ratio of soma spike interval to the axonal spike interval. The ratio is around unity, with a slight tendency to values >1 in longer branches. This deviation from the unity line is also observed in the upper right area (i.e., longer axons) of panel C.

Panel 2B exemplifies one somatic trace and its corresponding axonal trace. A *Mathematica* algorithm was developed to identify the peaks in both source and axonal traces, along with their timestamps. This process presents a challenge due to the noisy axonal voltage traces. To address this, the algorithm identified minimal amplitude peaks within a ±1 msec flanking the mean time of arrival (TOA). Figures 2C and D summarize analyses of 57 axonal loci and 4873 spike doublets from nine neurons across four distinct network preparations. In panel 2C, the axes, utilizing logarithmic scales, indicate time differences between the two spikes in the source (X-axis) and axon (Y-axis). Most of the points concentrate around the unity line. The histogram in panel 2D indicates that the ratio of soma spike interval to axonal spike interval is around unity, with a slight tendency to values >1 in longer branches. This deviation from the unity line is also observed in the upper right area (i.e., longer axons) of panel 2C. Altogether, the results indicate reliable conduction of intervals during spontaneous network activity.

Having observed that pairs of spontaneous, network-evoked spikes maintain their inter-spike intervals down the axonal tree, we turn to investigate the reliability of the axons when challenged by long series of repetitive external stimuli within a physiological range (1–10 Hz). To optimize the ability to interpret the measurements, we used synaptic pharmacological blockers (APV, CNQX, BIC; see Materials and Methods), thus significantly reducing the ongoing, spontaneous synchronous activity^26,29^. Neurons were stimulated either near the soma, or at the area identified as the likely axon initial segment based on the extracellularly generated neuronal footprint provided by the MaxLab Live software environment (MaxWell Biosystems AG). This allowed effective, reliable stimulation at voltages typically ranging from 150 or 200 mV. Balanced, positive-first biphasic voltage pulses were delivered locally at 1-3 neighboring electrodes at a frequency of 1, 3–4, or 10 Hz. Voltage stimuli evoked spikes time-locked to the downswing of the biphasic pulse.

When estimating the probability that spikes reach distal axonal arbors, the first task is to verify that the stimulus penetrates the axonal tree. Response failures are already common at the soma—even at physiological firing rates—and become more pronounced during stimulus trains^26,36,37^. We therefore define *responsiveness* at a given axonal site as the fraction of actual departures from the source. Specifically, we designate one or more *gate* electrodes (a cluster, when advantageous) and one or more downstream *target* electrodes that differ in path length and branch order from the gate (Figure 3). For each stimulus we mark every spike that leaves any gate electrode (logical union) and then record whether—and when—it is detected at eachb target. A failure at a target is scored only if no electrode in the target cluster registers an arrival (logical intersection).

**Figure 3:**
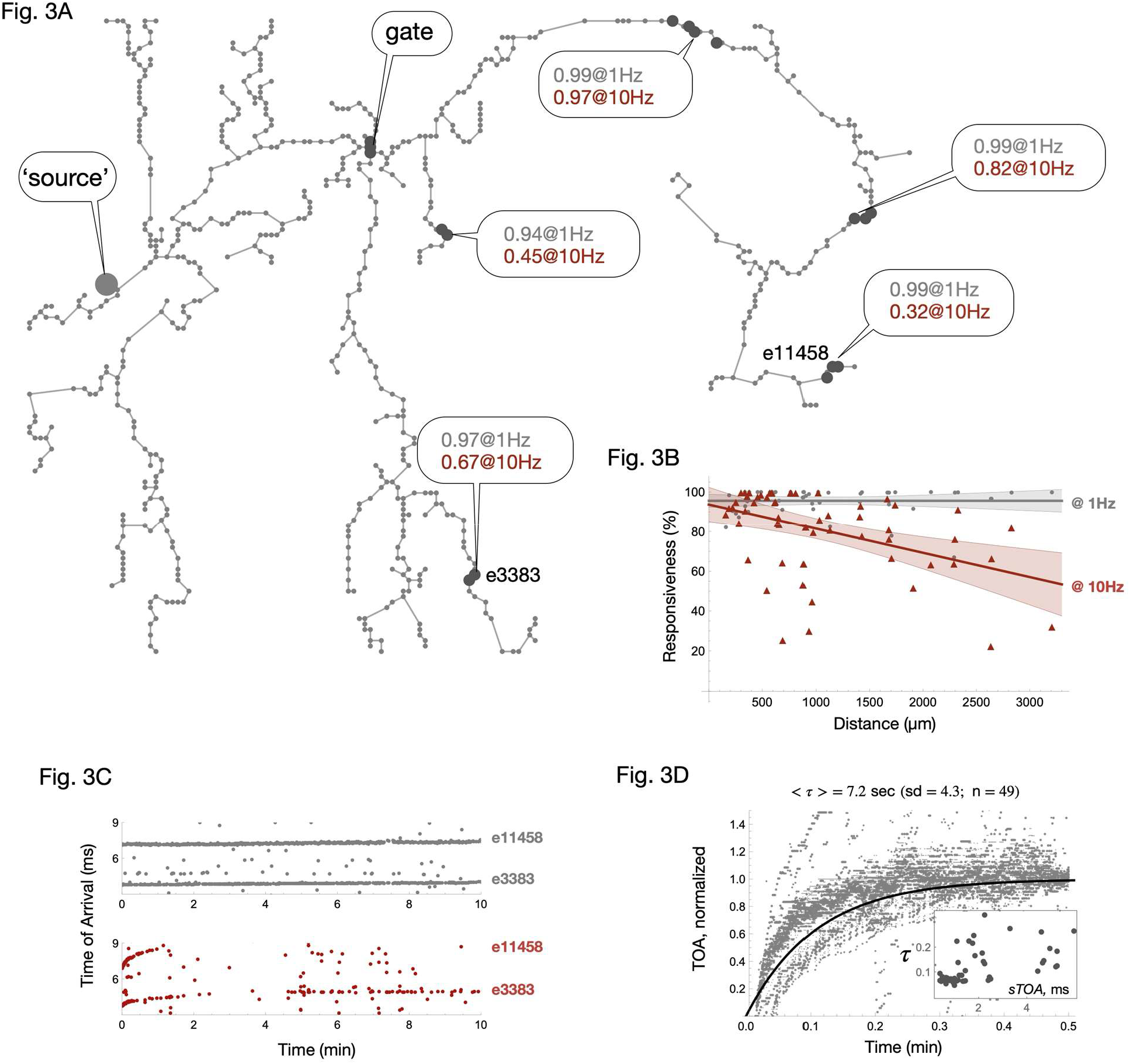
Response probabilities at different locations along the axonal arbor, responsiveness over time, and kinetics of TOA retardation. (**A**) Responsiveness at 1Hz (gray text) and at 10 Hz (red text) at different locations in the axonal tree for one example neuron. (**B**) Responsiveness versus soma distance at different locations along the axonal arbor for 1 and 10 Hz (1 Hz: 6 cells, 51 loci, 4 preparations; 10 Hz: 7 cells, 56 loci, 5 preparations). Correlation coefficients: r = –0.0004 at 1 Hz, r = –0.068 at 3 Hz (no meaningful relationship), but r = –0.45 at 10 Hz, indicating a strong negative correlation. (±95% confidence intervals for the mean response). (**C**) Response latencies and failure probabilities change during prolonged stimulation: responsiveness over time for a representative neuron (the one shown in Figure 3A). Dynamics of arrival times at two axonal electrode positions (depicted in Figure 3A) are shown. Time of arrivals (TOA, msec) to the depicted points along the axon vs. time (minutes) under stimulation at 1 Hz (top panel, gray), or at 10 Hz (bottom panel, red). (**D**) Kinetics of TOA retardation. Values were smoothed with a moving average (window size: 10 time points); scaled to the mean TOA of the last 10 points of the smoothed data (n = 49 traces, mean = 0.12 minutes, sd = 0.07, Median = 0.08 minutes, Range: Min 0.046, Max 0.33). Values are from 23 different axonal segments of 5 different neurons at 10 Hz. Black continuous line depicts an exponential function 1-e^-t/τ^, with the mean = 0.12 minutes) time constant of 49 traces. Inset: Time constant of TOA retardation as a function of spontaneous time of arrival (sTOA). There is a moderate-to-strong statistically significant positive correlation (r = 0.53, R^2^=0.28, p<0.001) between the two variables.

Traces are extracted from continuous recordings. Each trace spans 1–2 msec before a spike timestamp and 7–8 msec after it. For every electrode we compute a *quality index*—the ratio between the absolute minimum of its mean trace (maximal depolarization) and its root-mean-square (RMS) noise—and retain only electrodes that exceed a preset threshold. From these, we select a subset that are well separated spatially and positioned on both sides of major branch points. Axonal path lengths and branch orders are then derived from the neuron footprint electrodes with the algorithm described earlier: a shortest path tree is constructed, and each axonal electrode is annotated with its distance from the source and its branch order (tree depth). As an independent verification, the distance from the source is also estimated based on spike latencies measured during spontaneous activity recorded before synaptic blockers were applied. A correlation of 0.82 is observed between these two distance estimates.

The example neuron in Figure 3A reveals a pattern: At 1 Hz stimulation, spikes reach virtually every site in the axonal tree. At 10 Hz, by contrast, distance dependent propagation failures emerge. Figure 3B aggregates responsiveness versus soma distance for 1 and 10 Hz (1 Hz: 6 cells, 51 loci, 4 preparations; 10 Hz: 7 cells, 56 loci, 5 preparations). Correlation coefficients confirm the trend: r = –0.0004 at 1 Hz and r = –0.068 at 3 Hz (no meaningful relationship), but r = −0.45 at 10 Hz, indicating a strong negative correlation.

Although physiological theory—and classic large axon studies—implicates branch points, rather than path length itself, as the main sites of propagation failure^6,7,10,38,39^, our 10 Hz data do not permit a clear statistical partitioning of distance and rank effects. In our dataset, the two variables are highly correlated (r > 0.88), making it difficult to discern their individual contributions. We have attempted statistical methods to address the co-linearity (including Logistic Regression, Principal Component Analysis), but unfortunately, none have been successful.

Finally, we addressed *dynamics*: How do response latencies and failure probabilities change during prolonged series of stimuli? What is the temporal scale of such a change, and how does distance influence these changes? Our extended recordings, spanning up to 15 min, enable us to conduct this analysis. Previous studies had explored latency and failure dynamics, at higher stimulation frequencies (ranging from 40 to 140 Hz) and/or over brief periods of tens of seconds.^8,29,31^ Figure 3C plots the responsiveness over time for a representative neuron (the one shown in Figure 3A). At 10 Hz stimulation, spike latencies progressively lengthen within seconds, and failure rates start to appear, with both trends becoming more pronounced at greater soma distances (and, correspondingly, higher branch ranks). Figure 3D shows the kinetics of scaled TOA retardation from a range of axonal distant points. As indicated by the data of the inset, there is a moderate-to-strong statistically significant positive correlation (r = 0.53, R^2^=0.28, p<0.001) between the time constant of TOA retardation and the distance (represented in terms of spontaneous TOA). This effect might reflect the reduced probability of activation as the axonal distance increases (see Figure 3B).

### Interpretation and a Concluding Comment

Consistent with previous observations, spontaneous activity in large random cortical networks *in-vitro* unfolds over a wide range of time scales:^34,35^ inter-spike intervals are broadly distributed, exposing individual neurons’ firing irregularities enmeshed with network synchronous events that are not much different from *in-vivo* reported regimes.^34,35,40,41^ As shown above, under these conditions the axons produce a stable, accurate output that largely mirrors that of the soma. Axonal transmission remains reliable also during evoked activity at low-frequency (1–4 Hz) stimulation. When driven at 10 Hz for several seconds or more, delays lengthen and failures emerge, echoing a *critical frequency* reported earlier for these same cells.^26,36^ The effect intensifies with path length and (correspondingly) branch order.

To interpret these findings, we draw on extensive research, suggesting that activity reshapes the availability of voltage-gated sodium channels. Beyond the familiar fast inactivation that governs single spikes, these channels undergo *slow inactivation*, conferring a long memory of prior activity.^21–25,32,36,42–46^ NaV1.2 and by most accounts, also NaV1.6 channels— subtypes abundant in unmyelinated cortical axons^4,6,10,16^—display slow inactivation across timescales from tens of milliseconds to minutes.^47,48^ Time constants for recovery from slow inactivation scale with the duration of previous activity following power-law relations.^22–24,36,42,43,49^ This redistribution between available and non-available states can be approximated by a logistic equation,^36,42^ positioning the membrane near the boundary between excitable and non-excitable regimes.^36,37^ Spiking at this “edge of excitability” exhibits hallmarks of criticality—power-law relaxations, critical slowing, and adaptive scaling to input statistics—phenomena documented in somatic recordings of cortical neurons.^26,37,50^ Such critical-like behavior becomes especially consequential once spatial factors are introduced, given the vulnerability of conduction through volume inhomogeneities, thin distal branches, and—most prominently—branch points. Because a branch point increases the axial load, it is intrinsically the weakest link in that chain. The larger the geometrical load,^51^ or the smaller the sodium conductance availability, the lower the spike amplitude that reaches the daughter trunks. A long (tens of seconds) series of depolarizations at 10 Hz can reduce the availability of sodium conductance by ca. 20%,^22,42^ and position the membrane at the critical border for excitability where failures begin to appear.^36^

Simulations support this interpretation: expanding an axonal propagation model convolved with slow inactivation^32^ reveals increased latency and spike loss at branch points (Figure 4). At 10 Hz, a high Geometric Ratio (GR) (>1 and up to 2.5–reported in^e.g.,8,52,53^) leads to partial or complete propagation blocks at the branch or just before the branching point, or both. This phenomenon can arise from several biophysical and geometrical considerations: electrotonic discontinuity and local input impedance, active properties of non-uniformly distributed conductances, or even reflection block^2–4,6,7,10,12,13,20,32,54,55^.

**Figure 4:**
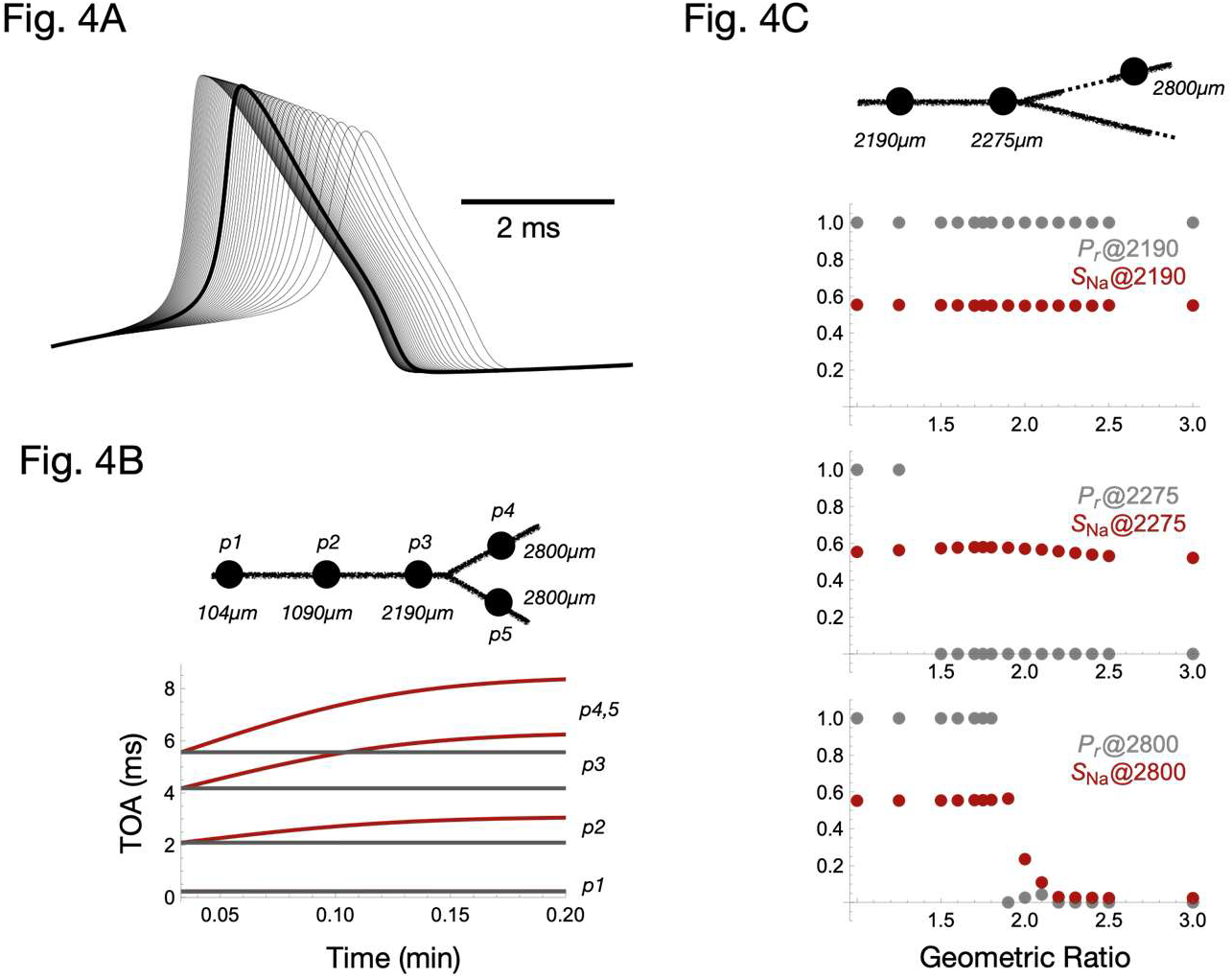
Sodium channel availability is a key determinant of propagation reliability in an axon model with slow inactivation. (**A**) Sensitivity of Hodgkin-Huxley model (single compartment) spike envelope to variation in sodium conductance availability (50–150 mS/cm^2^); thick black depicts “standard” 120 mS/cm^2^. (**B**) Latency pre and post the branching point under stimulation at 10 Hz with (red) and without (black) slow inactivation (Geometric Ratio GR = 1). (**C**) Average response probability (Pr) and fraction of sodium conductance in slow inactive state (SNa) pre and post the branching point under stimulation at 10 Hz at different GRs (1, 1.25, 1.5, 1.6, 1.7, 1.75, 1.8,1.9, 2, 2.1, 2.2, 2.3, 2.4, 2.5, 3). At GR 1.9 the branch point cannot overcome slow inactivation.

Altogether, the simulation suggests that reliable propagation at 10 Hz would require a maximal sodium conductance that is adequately high to safeguard against loss to slow inactivation; a demand for reliability at higher frequencies would increase the required maximum sodium conductance margin. Indeed, as demonstrated in several experimental studies^22,49,56,57^, repeated short activations of the membrane at frequencies as low as 2 Hz can induce inactivation at this level or higher.

In the present study, we report that axons maintain exceptional reliability, even under complex somatic firing patterns that include extremely short interspike intervals. Functionally, this supports the view of axons as robust conducting devices. This reliability, however, is not unlimited. As shown previously, artificially imposed high-rate pulsing stimuli can induce conduction failures; for cortical neurons, sub-second trains require frequencies above ∼80 Hz to do so, both *in vitro*^29^ and *in vivo*^8^. Extending this principle, we show that even 10 Hz stimulation can elicit failures if maintained for several seconds, with the probability of failure increasing with axonal distance and branching rank. We stress that such conditions arise primarily under experimental regimes that exceed typical physiological activity, though some specialized neurons may encounter sustained high-frequency firing *in vivo*. More plausibly, bursting activity, whose dynamics, as our analyses show, preserve reliable conduction, dominates cortical activity patterns. Thus, after all, axons may “only ax,”^1^ a hypothesis that remains to be tested under natural, ongoing *in vivo* conditions.

## Acknowledgments

We thank Danny Eytan, Naama Wolf, Itamar Kahn, Erez Braun, Omri Barak, Noam Ziv, Larry Abbott, and Ron Teichner for helpful comments. We thank Leonid Odessky and Yael Abuhatsera for invaluable technical assistance. This work was partially supported by research grants from the Israel Science Foundation (ISF 806/19 S.M.) and the Schaefer Scholars Program at Columbia University’s Vagelos College of Physicians and Surgeons (S.M., L.C.).

